# Late Embryogenesis Abundant (LEA) proteins confer water stress tolerance to mammalian somatic cells

**DOI:** 10.1101/704809

**Authors:** M Czernik, A Fidanza, FP Luongo, L Valbonetti, PA Scapolo, P Patrizio, P Loi

## Abstract

Late Embryogenesis Abundant (LEA) proteins are commonly found in organisms capable of undergoing reversible dehydration - “anhydrobiosis”. Here, we have produced three LEA proteins: pTag-RAB17-GFP-N, *Zea mays* dehydrin-1dhn, expressed in the nucleo-cytoplasm; pTag-WCOR410-RFP, *Tricum aestivum* cold acclimation protein *WCOR410*, binding to cellular membranes, and pTag-LEA-BFP, *Artemia franciscana* LEA protein group 3 that targets the mitochondria. Somatic cells transfected with three LEA proteins were subjected to desiccation under controlled conditions, followed by rehydration, viability assessment and membrane/mitochondria functional tests were performed. Results shown that LEA protect cells from desiccation injury. Cells expressed all LEA proteins shown very high percentage of viable cells (58%) after four hour of desiccation compare to un-transfected cells (1% cell alive). Plasmalemma, cytoskeleton and mitochondria appeared unaffected in LEA-expressing cells, confirming their protective action during the entire desiccation and rehydration process. Here, we show that natural xeroprotectants (LEA proteins) transiently expressed in somatic cells confer them desiccation tolerance.

## Introduction

Water is essential for life (Hand et al., 2007; Menze et al., 2009) yet many organisms are able to survive almost completely dehydrated (>99% of their body water is removed) (Crow et al., 1992). In nature, this phenomenon is known as “anhydrobiosis”, and is conserved across vegetal and animal phyla (Hincha et al., 1992). Anhydrobiosis allows seeds and small invertebrates to survive long time spans (decades/centuries) in the absence of water, thanks to the induced synthesis of sugars and various protein classes that can be collectively defined as “*xero-protectant*s” (Loi et al., 2013). Among them, Late Embryogenesis Abundant proteins (LEAp) are the best characterized and perhaps the most interesting (Marunde et al., 2013). LEA proteins were first discovered in cotton seeds more than 30 years ago (Dure et al., 1981) and were later also found in seeds and vegetative tissues of several other plants (Shih et al., 2008). A relatively recent survey, probably not updated, contains 769 LEAp entries from 196 organisms (Hunault and Jaspard, 2009). LEA proteins are highly hydrophilic and acquire random coils conformation in aqueous solution, property that has assigned them the definition of “intrinsically disordered” proteins (McCubbin et al., 1985). It is only during de-hydration that LEA proteins acquire their final conformations, primarily α helices, β sheets, and hairpin loops, and by doing so they bind to specific cellular/enzymatic substrates to be protected. The mechanism of “xero-protection” is not fully understood. Also lacking are data on the substrate binding mechanism. Data gained in model organisms have indicated that LEAp stabilisation occurs via several pathways: chaperon-like activity, protection of cell membranes, stabilisation of vitrified sugar glasses by increasing glass transition temperature (Tg), sequestration of divalent ions, and synergic interaction with other xero-protectants, such as trehalose (Li et al., 2012). The expression of LEAp, as well as the other xero-protectants, is triggered in anhydrobiotic organisms once water stress is sensed, leading to a progressive accumulation in various cellular compartments, such as mitochondria (Hand et al., 2011; Moore and Hand, 2016), nucleus (Wu et al., 2013) cytosol (hand et al., 2011), membranes (Tolleter et al., 2010), and endoplasmic reticulum. Clearly, several LEAp are required, along with other xero-protectants, to confer full protection from water stress. Therefore, if the strategy is to exploit LEAp for the induction of reversible drying in mammalian cells, it is necessary to know their specific targets and the mechanism of action. While the mechanism of action might be inferred from sequence analysis of the LEAp by bioinformatics, *in vivo* transfection assays of cells with the different LEAp are required to confirm their protective action, but also to exclude negative effects on cell homeostasis. After all, they are mainly vegetal proteins. An index paper published by Li’s group was the first to investigate the effects of LEAp expressed in mammalian cells subjected to rapid dehydration (Li et al., 2012). In that work, hepatoma cell line was stably transfected with a tetracycline (Tet)-inducible expression system coding for two LEAp naturally expressed in embryos of the brine shrimp *Artemia franciscana*, AfrLEA2 and AfrLEA3m, plus a trehalose transporter 1 (TRET1) (Li et al., 2012). The results showed that LEAp, one expressed in the cytoplasm, AfrLEA2, and the other AfrLEA3m in the mitochondria, together with trehalose, effectively protected the cells from the desiccation stress.

Our work extends Li’s findings. In addition to the LEAp AfrLEA3m, that targets the mitochondria, we have transfected primary cultures of fibroblasts with two additional LEA proteins: pTag-RAB17-GFP-N, *Zea mays* dehydrin-1dhn, expressed in the nucleo-cytoplasm; and pTag- WCOR410-RFP, *Tricum aestivum* cold acclimation protein *WCOR410*, that binds to the membranes. Our preliminary work on lyophilized, unprotected, somatic cells directly processed for scanning and electron microscopy showed massive damage at the membrane level [(Matzukawa, personal communication and (Iuso et al., 2012)]. Here reported results show that the individual LEAp protected the cells from desiccation with strongest effect when all three LEA were expressed adding new data on the induction of controlled drying in mammalian cells.

## Results

### Subcellular localization of pTag-RAB17-GFP, pTag-WCOR410-RFP, and pTag-LEA-BFP

Sheep fibroblasts were transiently transfected with pTag-RAB17-GFP, pTag-WCOR410-RFP, pTag-LEA-BFP with efficiency as follow: 44%, 26% and 24%, respectively. pTag-RAB17-GFP was localised to the cytoplasm and nucleus (Fig. 1A-C), pTag-WCOR410-RFP protein was observed only in the cytosol/membranes (Fig. 1G-S) and by localization with membrane dye clearly shown proper localization (Fig. 1M – S), while pTag-LEA-BFP was detected in the mitochondria (Fig.1T - Z). Fig. 1W-Z shows that pTag-LEA-BFP is targeted to the mitochondrial network as it co-localised with MitoTracker green (Fig. 1X, Z). Additionally, to confirm the proper localization of the LEA proteins, sheep fibroblasts were transfected with empty vectors EV-GFP, EV-RFP and EV-BFP as a control. Results showed that GFP, RFP and BFP alone had spread distribution throughout the cells (Fig. 1D-F: GFP; Fig. 1J-L: RFP; Fig. 1U: BFP). Expression of fusion LEA proteins in sheep fibroblast was also confirmed by immunoblotting analysis (Fig. 1Z’). Moreover, to our knowledge for the first time, we were able to express all tree LEA proteins (pTag-WCOR410-RFP (Fig. 2A); pTag-RAB17-GFP (Fig. 2B); pTag-LEA-BFP (Fig. 2C)) in the same somatic cell, although with low efficiency (11%). Additionally, in the MIX conditions we have also found cells that expressed only two LEA protein (12%) as well as cells with single LEAp: pTag-RAB17-GFP (21%), pTag-WCOR410-RFP (29%), or pTag-LEA-BFP (14%).

**Figure 1.**
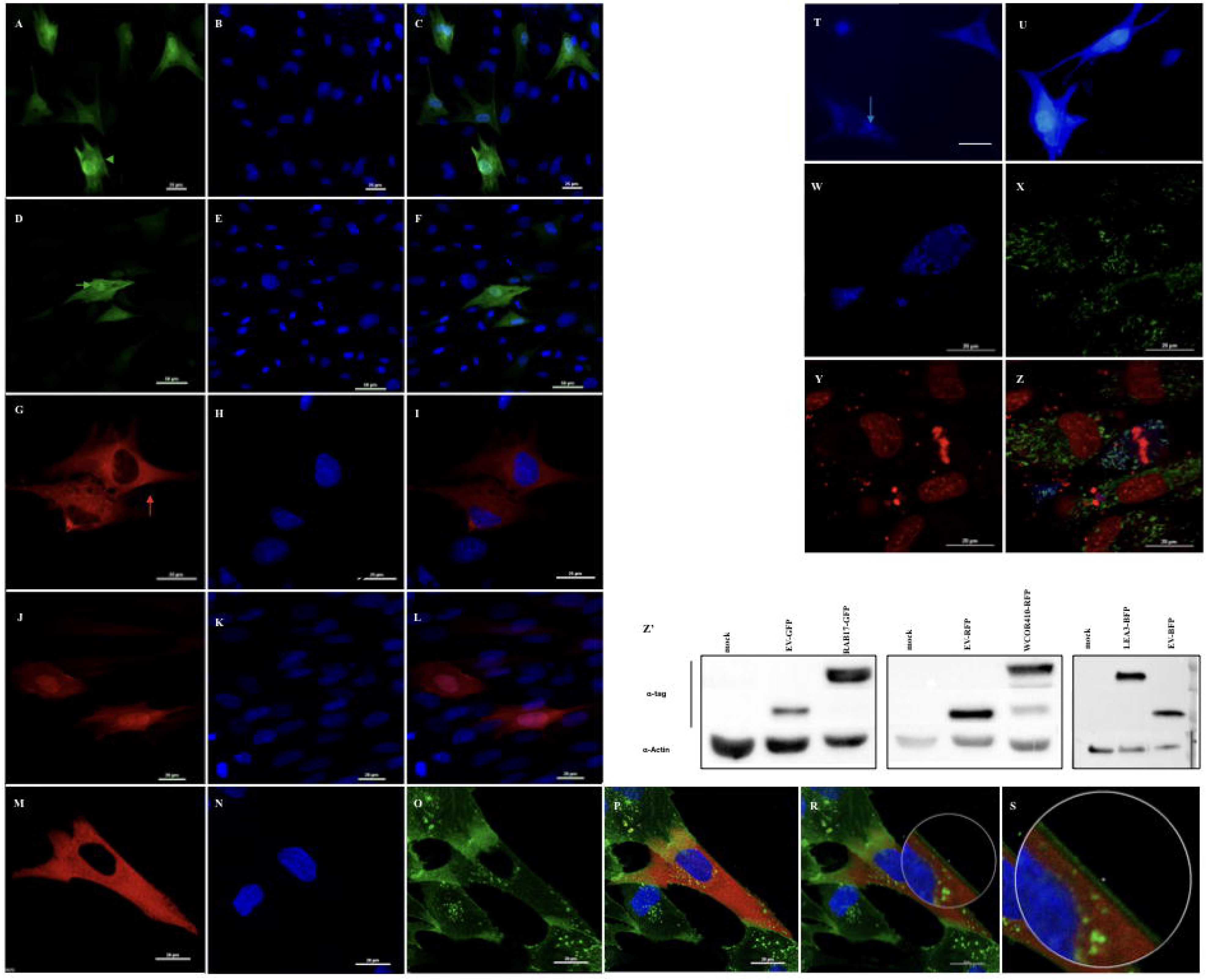
Subcellular localization of individual LEA proteins in sheep fibroblasts. (A-C) pTag-RAB17-GFP; (D – F) GFP-tag; (G - I) pTag-WCOR410-RFP; (J – L) RFP-tag; (M - S) pTag-WCOR410-RFP and membrane. (A) pTag-RAB17-GFP fusion protein shows cytoplasmic and nuclear localization (green arrows head, A); (B) nucleus stained with Hoechst 33342; (C) merge; (D) EV control expressing only GFP was spread throughout the cells; (E) nuclei stained with Hoechst 33342, (F) merge; (G) pTag-WCOR410-RFP fusion protein exhibit membrane localization (red arrow); (H) nuclei stained with Hoechst 33342; (I) merge; (J) RFP alone show spread localization in the cells; (K) nuclei stained with Hoechst 33342 (L) merge; (M) pTag-WCOR410-RFP fusion protein exhibit membrane localization (red arrow); (N) nuclei stained with Hoechst 33342; (O) membrane staning, (P) merge, (R, S) enlarge Fig. 1P; (T) pTag-LEA3-BFP shows mitochondrial localization, compared to (U) BFP alone that is spread throughout the cell; (W – Z) pTag-LEA3-BFP co-localizes with Mitotracker green; (W) expression of pTag-LEA3-BFP; (X) mitochondria stained with Mitotracker green dye; (Y) nuclei stained with Propidio Iode; (Z) merge; (Z’) expression of LEA proteins in sheep fibroblasts was verified by western blot.

**Figure 2.**
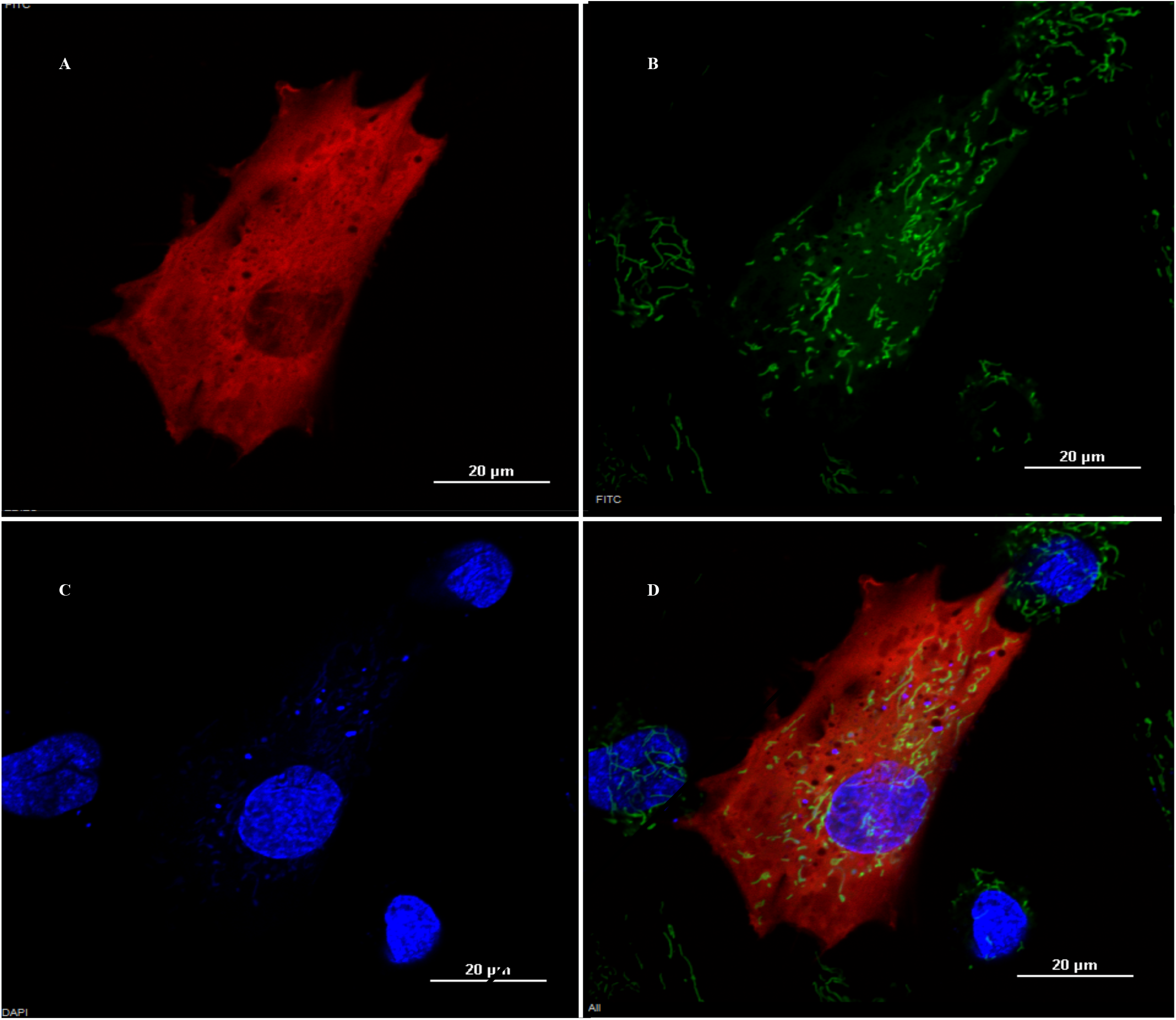
Subcellular localization of all three LEA proteins in sheep fibroblasts. (A) pTag-WCOR410-RFP fusion protein exhibited membrane localization; (B) pTag-RAB17-GFP fusion protein showed cytoplasmic and nuclear localization and mitochondira stained with Mitotracker green, (C) pTag-LEA-BFP showed mitochondrial localization as well as nucleus stained with Hoechst 33342 (D) merge. Red - RFP; green - GFP and Mitotracker green dye; blue - Hoechst 33342.

### Enhanced resistance to desiccation stress

Sheep fibroblast transfected with LEA proteins, as well as not transfected controls (CTR), were air dried at 16°C for up to 4h. Every 60 min, cells viability was assessed on sub-samples using trypan blue exclusion. The results showed that 1h of air drying did not affect viability of sheep fibroblasts (Fig. 3A), with more than 80% of cells expressing any of the LEAp singularly or together and 60% of CTR maintaining viability (Fig. 3A). One hour later small negative effect of desiccation on cells viability start to be observed but any statistic differences between LEAs and CTR groups were observed. Drastic differences were observed 1h later (3h after initiation of desiccation). The numbers of viable cells expressing a single LEA protein were significantly higher than control group (RAB-17: 16%, WCORB410: 13%, LEA3: 12%, CTR: 2%). By 4h after desiccation begun, only few sheep fibroblast in the CTR were still alive (less than 1 %) while LEA proteins were able to protect the somatic cells, as indicated by their viability (Rab17: 8%, WCOR410: 5%, LEA3: 2.3%) (Fig. 3A), with the stronger effect observed when all three LEA proteins were co-transfected together (LEA-MIX). In this group, 40% of LEA-MIX transfected cells were still alive after 3h of air drying, compared to 2% of the CTR group; 23% of MIX cells were viable after 4h of drying while in the control group viability dropped to under 1% (Fig. 3A).

**Figure 3.**
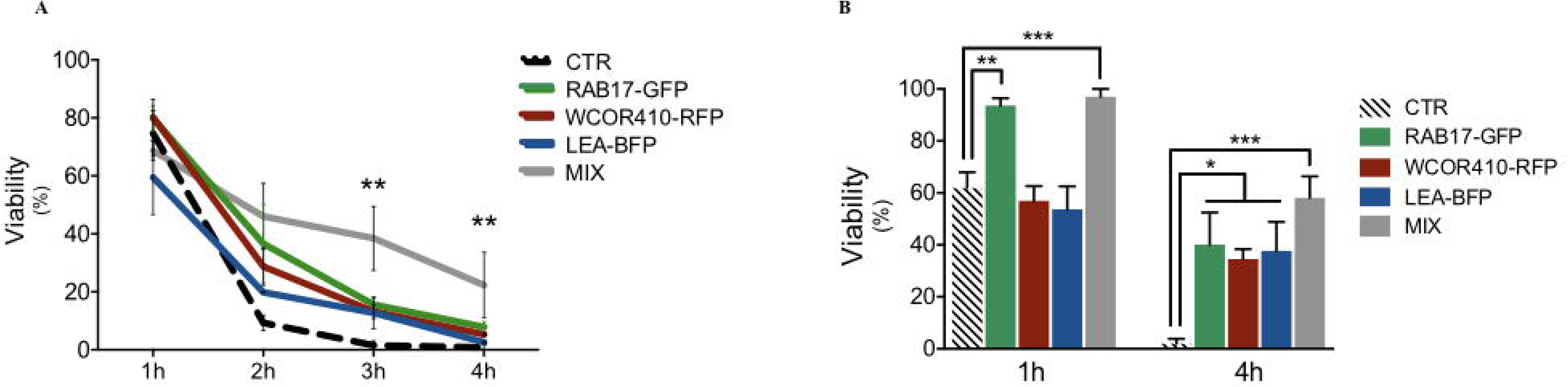
Enhanced resistance to desiccation stress. (A) Sheep fibroblast transfected with single LEA protein or with all three LEA proteins (here called MIX). Viability of the cells was controlled every hour for up to four hours using Tripan Blue staining; ** means value P=0.0052 CTR vs MIX (3 and 4 hours) (B) Sheep fibroblast transfected with single LEA protein or with MIX. Viability of the cells was controlled at two time points (after 1h and 4h of controlled drying); **means P=0.03 CTR vs RAB17 after 1h of desiccation; *** means P=0.0046 CTR vs MIX after 1h and 4h of desiccation; *p<0,05 CTR vs single LEAs 4h post desiccation.

To verify viability every hour, we were forced to remove the cells from the drying chamber, and by doing so we exposed them to uncontrolled variation in both humidity and temperature. This could have affected the late time points. For this reason, we decided to carry on the viability tests at two time points: 1h and 4hs post desiccation. Accordingly, stronger difference in cells viability was observed when cells were not exposed to condition changes. Indeed, after 4h of desiccation, statistically significant difference between cells expressing single as well as all three LEA together (MIX) and the control group were observed (RAB17: 40%, WCOR410: 34%, LEA: 37%, MIX: 58%, CTR: 2%) (Fig. 3B).

### LEA proteins preserve proliferation capacity following desiccation

Transfection of a single LEA protein resulted in a higher number of cells attached to culture dishes (pTag-RAB17-GFP: 55 cells/field; pTag-WCOR410-RFP 62 cells/field; pTag-LEA-BFP: 37 cells/field; LEA-MIX: 86 cells/field) than in cells desiccated without LEA, CTR-D (non-transfected and desiccated cells) (25 cells/field), (Fig.4B). Proliferation rate was 32% with pTag-RAB17-GFP; 31% with pTag-WCOR410-RFP; and 25% with pTag-LEA-BFP (Fig. 4A). In the LEA-MIX group, cell proliferation rate was at levels comparable to the non-desiccated controls (CTR) (48% vs 51%, respectively).

**Figure 4.**
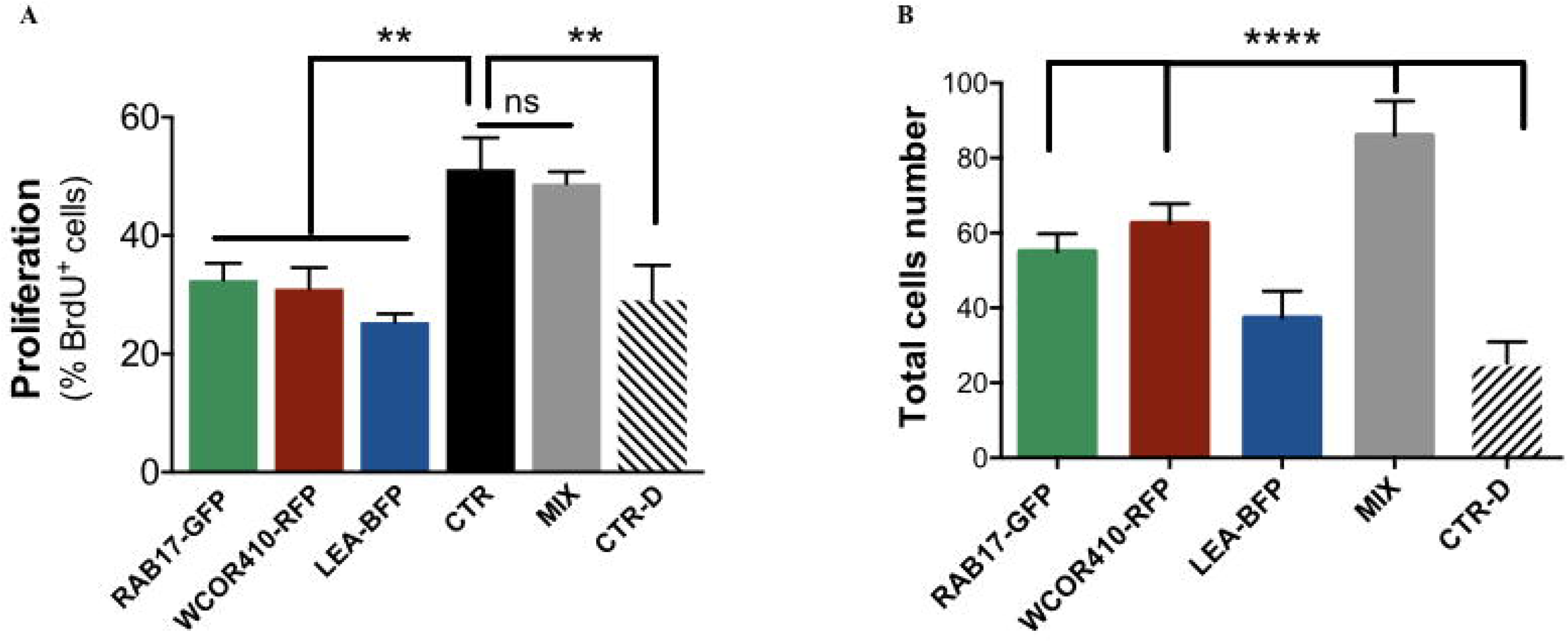
LEA proteins preserve proliferation capacity following desiccation. (A) Proliferation was measured by counting of cells that had incorporated BrdU in the nuclei; ** p<0,005; *** p<0,0005; (B) Attached cells count after desiccation, re-hydration and culture for 24h in incubator [(transfected with single LEA proteins: pTag-RAB17-GFP, pTag-WCOR410-RFP, pTag-LEA-BFP or all together (MIX)];

### Cellular integrity of mammalian cells after desiccation

Somatic cells transfected with LEA proteins and controls (non-transfected and desiccated – CTR-D), were cultured for an additional 24h after desiccation and rehydration. Non-transfected and non-desiccated cells were used as a positive control (CTR). Results showed that LEA proteins protected cellular organelles from desiccation injury (Fig. 5). Cells expressing pTag-WCOR410-RFP (Fig. 5A-H) and pTag-RAB17-GFP (Fig. 5I-R) did not show any post-desiccation damages of the cytoskeleton as shown by F-actin staining (Fig. 5A, O). Normal actin filaments, spanning the entire cells, were observed in sheep fibroblast expressing these LEA proteins, compared to the CTR group (Fig. 5T). CTR-D cells were less organised and showed fragmented cytoskeleton (Fig. 5Y). Importantly, LEA proteins, and particularly pTag-LEA-BFP, protected the mitochondria in transfected cells (Fig. 6A-D), where high numbers of active mitochondria localized in perinuclear position (Fig. 6A), were observed, similar to the CTR group (Fig. 6E-H). On the other hand, mitochondrial activity was very poor in CTR-D group (Fig. 6I-L) where the organelles were localized peripherally, displaying high fragmentation, and were much less metabolically (Fig. 6I).

**Figure 5.**
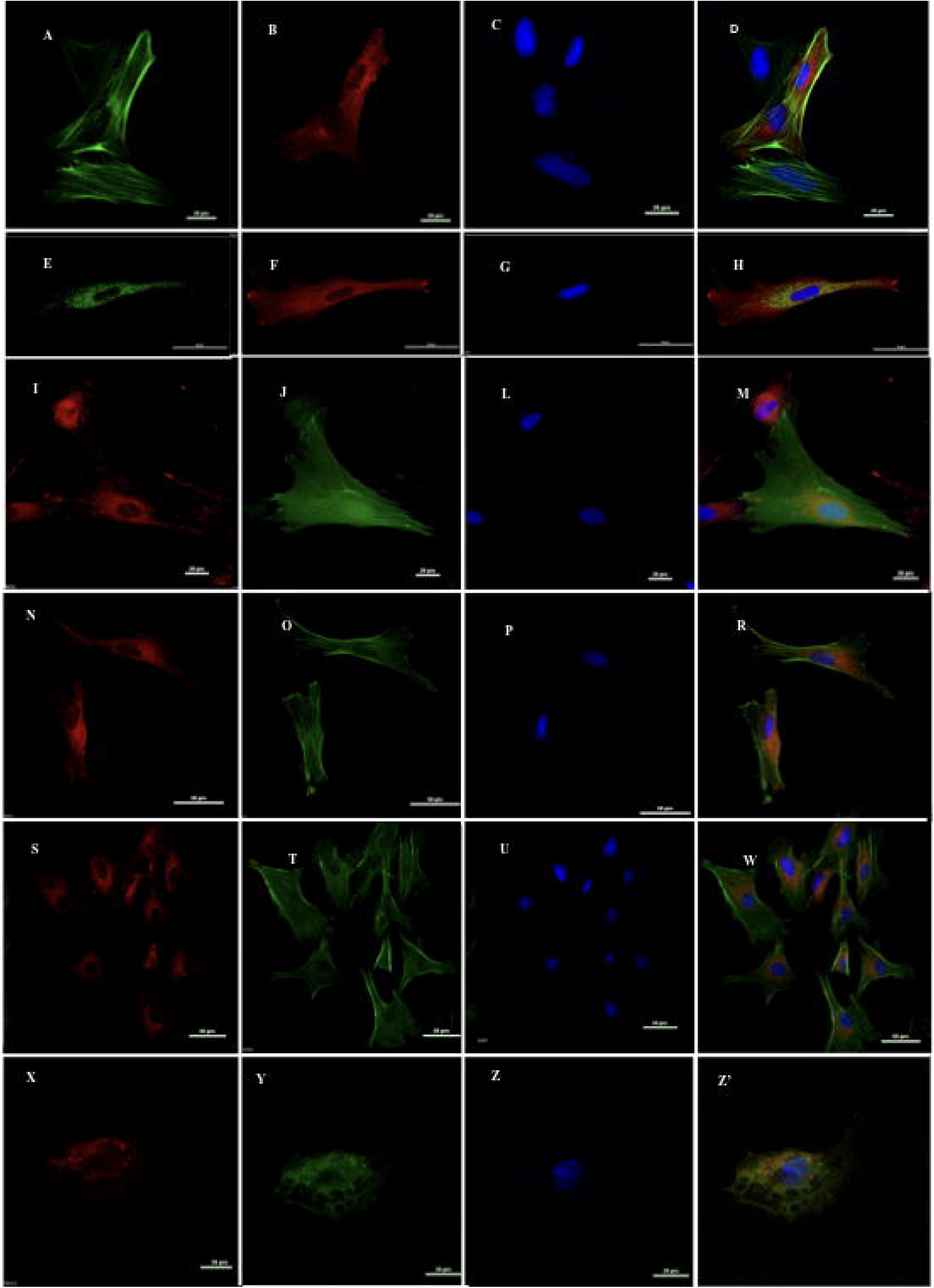
Integrity of mammalian cells after desiccation. (A-H) sheep fibroblasts expressing pTag-RAB17-GFP; (I-R) sheep fibroblasts expressing pTag-WCOR410-RFP; (S–W) non desiccated control sheep fibroblasts (CTR); (X-Z’) non transfected sheep fibroblasts, after desiccation (CTR-D); (A, O, T, Y) cytoskeleton, stained with Phalloidin green; (I, N, S, X) mitochondria, stained with Mitotracker red dye; (E) mitochondria, stained with Mitotracker green dye; (C, G, L, P, U, Z) nucleus, stained with Hoechst 33342; (D, H, M, R, W, Z’) merge;

**Figure 6.**
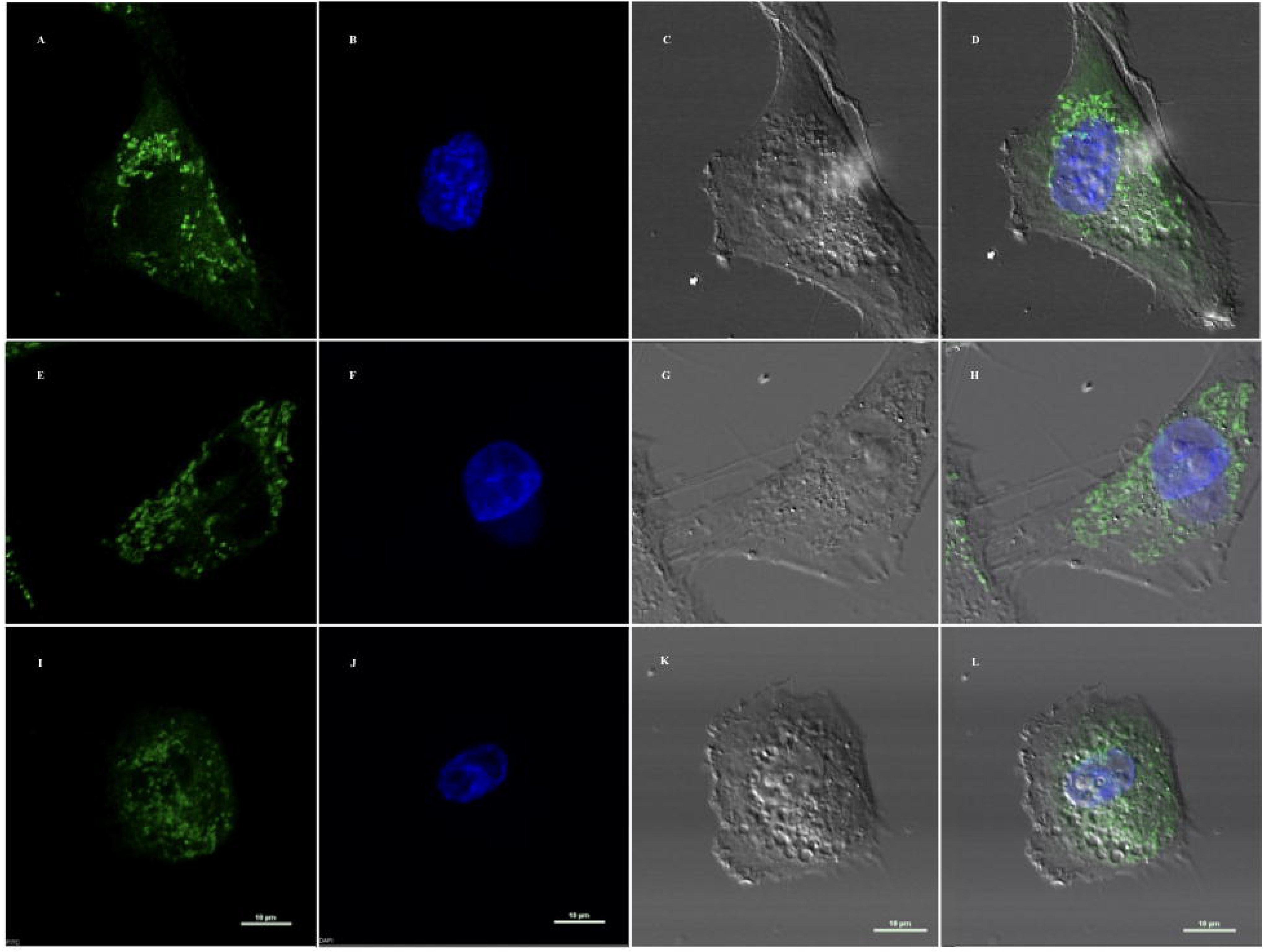
pTag-LEA-BFP protects the mitochondria from desiccation damages. (A – D) sheep fibroblasts expressing pTag-LEA-BFP; (E – H) control non-desiccated sheep fibroblast s(CTR); (I – L) control fibroblasts subjected to desiccation (CTR-D), green - mitochondria stained with Mitotracker green dye; blue - nucleus stained with Hoechst 33342.

## Discussion

The natural capacity of simple organisms to survive in a dehydrated state has long been exploited by humankind, with lyophilization as the method of choice for the long-term storage of bacteria and yeast (Fonseca et al., 2015). Some attempts were subsequently conducted to freeze dry non-nucleated mammalian cells, like platelets and red blood cells, with partial success (Crowe et al., 2003). It was the report of the maintenance of nuclear viability in lyophilized spermatozoa (Wakayama and Yanagimachi, 1998) that sparked the interest in dry stabilization of mammalian cells. The low water content and the highly condensed DNA make spermatozoa “easy” to dry, while in contrast other mammalian cells are not tolerant to dehydration and invariably die. No genes related to LEA family have been identified in the mammalian genomes sequences so far, beside one short protein conferring mitochondrial protection (Hall et al., 2011). Therefore, the only possible way to confer cell desiccation tolerance is to provide them with suitable xero-protectants.

The first studies reporting on the possibility of drying somatic cells were published by Guo et al., (2000) and Eroglu et al., (2000). These authors showed that it was possible to desiccate and store human fibroblasts for up to 5 days while maintaining viability upon rehydration. Their method made use of the protective effects of trehalose, a disaccharide associated with organisms withstanding desiccation (Leslie et al., 1995; Welsh and Herbert, 1999). Trehalose was produced in cells previously infected with an adenoviral vector expressing the trehalose biosynthetic genes, otsA and otsB, followed by air-drying and storage at room temperature.

Here we followed this general strategy, but LEA proteins were used as xero-protectants. Our work builds on a recent paper where desiccation tolerance was induced in hepatoma cell line expressing Tet-inducible expression system coding for two LEA proteins of the brine shrimp *Artemia franciscana*, AfrLEA2 and AfrLEA3m, and a trehalose transporter 1 (TRET1). Of the two LEA proteins, AfrLEA2 accumulated in the cytoplasm, and AfrLEA3m selectively targeted the mitochondria. Here we maintained the latter LEA protein, but we selected two different additional ones. In our previous work on lyophilization and nuclear transfer of lyophilized cells, we found high level of DNA damage in the resulting pronuclei (Iuso et al., 2012). Therefore, we elected to use pTag-RAB17-GFP-N, *Zea mays* dehydrin-1dhn, that is expressed not only in the cytoplasm as AfrLEA2 does, but also in the nucleoplasm, to protect the DNA as well. The third LEA protein that we utilized was pTag-WCOR410-RFP *Tricum aestivum* cold acclimation protein *WCOR410*, that binds specifically to the membranes. This latest LEA proteins was included following our observation of a massive membrane damage in lyophilized fibroblasts processed for scanning electron microscopy (SEM) without re-hydration (Matsukawa, unpublished) and for transmission electron microscopy (TEM) after rehydration (Iuso et al., 2012).

The proportion of fibroblasts expressing single LEA proteins was around 30%, with no significant differences between the three vectors. Transfection efficiency of all three LEA expression vectors was much lower, about 11%. As a result, the data on cell survival were underestimated because desiccated samples contained both expressing and non-expressing cells. Transgenic cell lines constitutively expressing xero-protectants, as accomplished in Li and co-workers (Li et al., 2012), provide indeed more precise and objective findings, but such approach cannot be an option for practical use.

The subcellular localization of all three LEA proteins matched the expectations (Fig. 1 and 2), confirming an earlier report (Li et al., 2012). No adverse effects on cell viability were observed in the LEAp expressing cells.

The LEA proteins exerted protection against water deprivation, with no major differences between them. LEA pTag-RAB17 expression appeared to be more beneficial over the other two, probably owing to its ubiquitous expression in all cell compartments (Fig. 3 and 4), and also because of its higher transfection efficiency compared to the other two [(44% vs 26% (WCOR410-RFP) and 24% (LEA3-BFP)]. Clearly, cells expressing all three LEA proteins showed the best survival rate, particularly in the experiment with only two time point controls (Fig. 3B). Expression rate of all LEA proteins in the same cells was low (11%) but presence of other LEAp combinations (single LEAp expression, and combinations of two LEAp in the same cell) increased the protective effects in the MIX group.

The proliferation assays after the water stress further demonstrated the beneficial effects of LEA proteins, basically in the cell functions explored - mitochondrial function and distribution, and F-actin (cytoskeleton). Again, no major differences between cells expressing individual LEA protein were detected (Fig. 5), while those expressing the three LEA proteins showed remarkable growth performances, comparable to control, unstressed cells (CTR) (Fig. 4A). Viability was further supported by the normal number and distribution of the mitochondria in growing cells, as well as the normal organization of polarized F-actin across the cells (Fig. 5). In contrary, unprotected cells displayed abnormal mitochondrial distribution (Fig. 6I) and disordered F-actin scattering across the cells (Fig. 5Y), suggesting that time is needed to recover from desiccation damages.

As learned from anhydrobiotic models, desiccation tolerance is conferred through a combined action of several xero-protectants, including LEA proteins. Our work using a peculiar combination of three LEAp expressing plasmid vectors transferred into sheep fibroblasts, reconfirmed their predicted localizations (Fig. 1), and convincingly demonstrated their protective effects during dehydration, the recovery on rehydration and continued growth following *in vitro* culture. Our data ameliorate our knowledge on the induction of reversible drying in mammalian nucleated cells for their long-term stabilization in an anhydrous state as an alternative biobanking approach.

## Materials and methods

### Construction of LEAp plasmids

Coding Sequences (CDS) for the LEA protein were produced by gene synthesis (Dundee Cell Products, USA): *RAB17* (~0.5 Kb; *Zea mays* dehydrin-1dhn, GenBank NM_001111949.1), *WCOR410* (~0.8 Kb; *Tricum aestivum* cold acclimation protein *WCOR410*; GenBank L29152.1) and LEA (~0.9 Kb; *Artemia franciscana* LEAp group 3; GenBank FJ592175.1). *RAB17*, *WCOR410* and *LEA* were subcloned into pET-15b (Novagen, Rome, Italy) under T7 promoter. Subsequently, CDS were amplified using AccuPrime Pfx DNA polymerase (ThermoFisher), and inserted using EcoRI/HindIII into the pTag-GFP-N, pTag-RFP-N to obtain pRAB17-GFP and pWCOR410-RFP, and SacI/PstI for the pTag-BFP-N to obtain pLEA-BFP (all plasmid backbones were from Evrogen, Milano, Italy). Correct clones were confirmed by Sanger sequencing using ABI PRISM 3100 (Applied Biosystem).

### Transfection

Sheep adult fibroblast (SAF) were derived from ear biopsy of three female Sarda breed sheep (2 years old). Animal work (skin biopsy) has been approved by the Italian Ministry of Health, upon the presentation of the research description prepared by the ethics committee of the Istituto Zooprofilattico Sperimentale di Teramo (Prot. 944F0.1 del 04/11/2016). The number of the authorization granted by the Italian Ministry of Health is n° 200/2017-PR. We confirm that all methods were performed in accordance with the relevant guidelines and regulations.

SAFs (between second and eighth passage) were cultured in DMEM (GIBCO) containing 2 nM glutamine, 3.7 g/L NaHCO3 and 0.5% gentamicin supplemented with 10% Fetal Bovine Serum (FBS).

Transfection of sheep adult fibroblasts was adapted from Czernik and colleagues (Czernik et al., 2016) with small modifications. Approximately 10^6^ cells were plated in 3,56 cm dishes and cultured in Minimal Essential Medium (MEM) + 10% foetal bovina serum (FBS) (Gibco, Milan, Italy) for 24h. After 24h cells were transfected using Lipofectamine 2000 kit (Invitrogen, Milan Italy), according to the manufacturer’s protocol, using 3μg of pTag-RAB17-GFP-N, pTag-WCOR410-RFP-N, pTag-LEA-BFP, -individually or in combination (by 2μg of each) (called MIX). Additionally, empty vectors: pTags-GFP-N, pTags-RFP-N, pTags-BFP-N were used as a control (called EV-GFP, EV-RFP, EV-BFP, respectively). After transfection cells were incubated in a humidified atmosphere 5% CO_2_/95% air at 37°C. All experiments were done 24h post-transfection. Efficiency of the transfection of all experiments were normalised normalized by using an internal control,

### Localization of LEA proteins in somatic cells

Sheep fibroblasts transfected with pTag-RAB17-GFP, pTag-WCOR410-RFP, pTag-LEA-BFP individually or in combination, as well as empty vector controls, EV-GFP, EV-RFP, EV-BFP, were fixed with 4% paraformaldehyde (PFA) for 20 min at room temperature (RT). After subsequent wash with PBS, cells were counterstained for 10min with 5μg/mL Hoechst 33342 (pTag-RAB17-GFP and pTag-WCOR410-RFP cells) or 0.5μg/mL Propidium Iodide (PI) (pTag-LEA-BFP expressed cells). Then, cells were mounted on slides with Fluoromount™ aqueous mounting medium (Sigma, Milan, Italy) and localization of the proteins was analysed with using Nikon Ar1 laser confocal scanning microscope (Nikon Eclipse Ti-E) equipped with the NIS-Element 4.40 software.

### Western Blot

Proteins were isolated from sheep fibroblasts transfected with pTag-RAB17-GFP, pTag-WCOR410-RFP, pTag-LEA-BFP, EV-GFP, EV-RFP, EV-BFP and from the mock transfection control, by incubation overnight with lysis buffer I (20 mM Tris, 150 Mm NaCl_2_, 1% NP-40) at 4°C. Then protein extracts were re-suspended at 1:1 ratio in lysis buffer II (20 Mm Tris, 150 Mm NaCl_2_). Protein concentration was assessed using the BCA Protein Assay Kit (Thermo-Fisher, Milan, Italy) according to the manufacture protocol. For each sample 50 μg of protein were incubated at 95°C for 10 min and then loaded into a gradient (4-15%) western blot gel (mini-protein TGX gel, Bio-Rad, Milan, Italy). Proteins were transferred onto a 0.45 μm nitrocellulose membrane (Bio-Rad, Milan, Italy) at 4°C for 2h at 200 mA. After transfer, the nitrocellulose membrane was blocked with 5% non-fat dry milk in 0.1% Tween-20 PBS (PBST) for 1h at RT. Membranes were incubated overnight with rabbit anti-tagRFP (which recognise also tagBFP) or rabbit anti-tag(CGY)FP primary antibodies (both from Evrogen, Milan, Italy) at 1:5000 in PBST with 0.5% non-fat dry milk. Then, membranes were washed three times for 15 min with PBST and incubated with the secondary antibody donkey anti-rabbit- IgG-HRP (sc-2317, Santa Cruz Biotechnology, USA) at 1:10000 for 1h at RT. Final detection was performed using enhanced chemiluminescence (ECL) Western Blotting Substrate (Amersham-Pharmacia, Piscataway, NJ, USA) and image acquisition that was carried out using the ChemiDoc System (Bio-Rad, Milan, Italy). Western blot analysis were repeated 4 times.

### Desiccation of sheep fibroblasts expressing LEA proteins, cell viability, and residual water

Sheep fibroblasts transfected with pTag-RAB17-GFP, pTag-WCOR410-RFP, pTag-LEA-BFP, individually or in combination, were detached with Trypsin-EDTA (0.25%) and pelleted by spinning them for 5 min at 1200 rpm (Eppendorf Centrifuge 5804). Cells were then re-suspended to 10^5^ cells/mL in desiccation medium (50 mM Hepes and 500 mM trehalose in PBS) and drops of 10μL were placed on a plastic, cover dish. Cells were then air-dried at 16°C for 1,2 3 and 4h. After desiccation, cells were rehydrated by adding 50 μL of the same desiccation medium and incubated for 5 min at RT. Viability and number of alive cells were evaluated using Trypan Blue staining. Un-transfected cells were used as a control (CTR). Residual water was assessed by weighing first the empty cover dish, then weighing the samples before drying and again after drying. This gave us a value of residual water per dry weight. Desiccation and residual water assessment was done 15 times.

### Cell proliferation assay

After desiccation, the fibroblasts were transferred into culture medium (MEM + 10% FBS) and cultured in a humidified atmosphere (5% CO_2_/95% air at 37°C) for 24 h. Cell proliferation was assessed by indirect immunocytochemistry detection of 5-bromo-2’-deoxyuridine (BrdU), a thymidine analogy incorporated during the S-phase of the cell cycle. Briefly, cells were cultured with 100 μM BrdU for 6 hours before the end of culture, fixed in cold 100% methanol for 20 min, and permeabilised with 0.1% Triton-X-100 in PBS for 15 min at RT. Next, cells were treated with 4N HCl at RT for 30 min and incubated with mouse anti-BrdU at 1:100 (B2531, Sigma, Milan, Italy) in blocking solution (0.1% BSA in PBS) over-night at 4°C, overnight. Cells were then incubated with rabbit anti-mouse IgG-FITC polyclonal antibody at 1:500 (F9137, Sigma, Milan, Italy) in blocking solution at RT for 2h and counterstained with 0.5μg/mL PI at RT for 5 min. Between passages, cells were washed twice with PBS at RT for 5 min. Proliferation assay were repeated 5 times and at every repeat 10 different field/sample were photographed. The number of proliferative cells (expressed BrdU) vs. total cells number were automatically counted using Nikon Ar1 laser confocal scanning microscope (Nikon Eclipse Ti-E) equipped with the NIS-Element 4.40 software.

### Mitochondrial and cytoskeleton staining

After desiccation, the fibroblasts were transferred into the culture medium (MEM + 10% FBS) and cultured in a humidified atmosphere (5% CO_2_/95% air at 37°C) for 24h. The cells were then incubated with 1 μM of Mitotracker green FM (Invitrogen, Molecular Probes, Milan, Italy) for mitochondrial staining or 1 μM of Phalloidin green (Thermofisher, Milan, Italy) for F-actin (the cytoskeleton) staining, both in serum-free MEM for 30 min at 38.5°C. Then, cells were washed twice with PBS, fixed with 4% paraformaldehyde and counterstained with 5μg/mL of Hoechst 33342. All slides were examined by confocal microscopy (Nikon Eclipse Ti-E) using NIS-Elements Confocal software (Nikon).

### Statistical analysis

One-way ANOVA and Fisher exact test were used to compare live cells at different durations of desiccation. Data reported in this paper are the mean (±SEM) for each group. The level of significance was set at *P* < 0.05. Statistical analyses were performed using GraphPad Prism for Windows (Version 6.01, GraphPad Software, Inc, CA, USA).

## Acknowledgements

The project has received funding from the European Union’s Horizon 2020 Research and Innovation Programme under the Marie Skłodowska-Curie grant agreement No 734434 (DRYNET) and by Narodowe Centrum Nauki (NCN) GA 2016/21/D/NZ3/02610 to MCz.

## Author Contributions

M.C, A.F conceived and designed the research; M.C, A.F, FP.L performed the experiments; M.C, A.F, FP. L, L.V analyzed the data; AP.S and PP edit the paper M.C, A.F and P.L wrote the paper. All the authors discussed the results and contributed to the writing edited and reviewed the manuscript.

